# Predicting the dynamic interaction of intrinsically disordered proteins

**DOI:** 10.1101/2023.12.04.569847

**Authors:** Yuchuan Zheng, Qixiu Li, Maria I. Freiberger, Haoyu Song, Guorong Hu, Moxin Zhang, Ruoxu Gu, Jingyuan Li

## Abstract

Intrinsically disordered proteins (IDPs) participate in various biological processes. Interactions involving IDPs are usually dynamic and affected by their inherent conformation fluctuations. Comprehensive characterization of these interactions based on current techniques is challenging. Here, we present GSALIDP, a GraphSAGE-LSTM Network to capture the dynamic nature of IDP-involved interactions and predict their behaviors. This framework models multiple conformations of IDP as a dynamic graph which can effectively describe the fluctuation of its flexible conformation. The dynamic interaction between IDPs is studied and the datasets of IDP conformations and their interactions are obtained through atomistic molecular dynamic (MD) simulations. Residues of IDP are encoded through a series of features, including their frustration. GSALIDP can effectively predict the interaction sites of IDP and the contact residue pairs between IDPs. Its performance in predicting IDP interaction is on par with or even better than the conventional models in predicting the interaction of structural proteins. To the best of our knowledge, this is the first model to extend the protein interaction prediction to IDP-involved interactions.

## Introduction

Protein-protein interactions (PPIs) play a central role in all biological processes ^1–3^. Conventionally, PPIs are about structural proteins ^4–8^. However, there are numerous proteins that lack stable structures and are denoted as intrinsically disordered proteins (IDPs) ^9–11^. IDPs are widely involved in interactions with other proteins. These interactions are usually dynamic which enable IDPs to mediate malleable molecular recognition ^12^. Modes of IDP-involved interactions include fuzzy binding, coupled folding and binding, and disordered complex formation ^12–18^. These dynamic interaction modes are distinct from conventional PPIs. And they are essentially related to biological processes such as transcription regulation, cell signaling, and the formation and function of membraneless organelles ^19–23^.

There are growing numbers of studies to depict IDP-involved interactions ^3,14,15,18,21,24–28^. For example, experiments based on FRET and NMR were conducted to demonstrate the binding affinity and sequence tendency of interactions^18,26,29^. Despite these progresses, direct information about the dynamic nature of IDP-involved interactions and its relevance to the malleable binding still remain elusive which is largely due to the limitation of spatial or temporal resolutions ^30^. Simulations were widely applied to study the behavior of IDP-involved interactions ^15,24,26^. As revealed in our previous molecular dynamic (MD) simulation the interactions between IDPs are coupled with their conformation fluctuations ^15^. Reliable depiction of such dynamic interaction requires sufficient simulations of this complex, high-dimensional system. Simulation studies thus face considerable challenges due to the formidable computational costs. The state-of-the-art machine-learning methods have the potential to effectively investigate IDP-involved interactions because of their efficiency in recognizing the patterns of high-dimensional data ^31^.

Machine-learning methods have been widely used in the prediction of the interaction behaviors of structural proteins ^32–44^. These PPI prediction methods can be classified into two classes, the sequence-based methods that only require protein sequences, and the structure-based methods which also utilize protein structures. However, direct application of these methods in IDP-involved interactions is very difficult. IDPs have low sequence conservation and high proportions of low-complexity sequence domains (LCDs). Such sequence features provide challenges for sequence-based models in extracting information from IDP sequences ^45^. In addition, due to the limitation of experimental resolution, the structures of both monomeric IDP and IDP-involved complex are largely unavailable. Moreover, structure-based models are designed to make predictions based on the single stable structure, largely ignoring the conformation fluctuation of protein. The structure-based methods for predicting IDP-involved interactions are largely absent. And the development of prediction methods for IDP-involved interactions should effectively consider their dynamic natures such as their couplings with conformation fluctuations of IDPs.

In this work, we develop a GraphSAGE-LSTM Network (noted as GSALIDP) for predicting the behavior of IDP-involved interactions. By modeling multiple IDP conformations as a dynamic graph, this framework can effectively characterize the fluctuation of its flexible conformation. Here, the dynamic interaction between IDPs is considered and the datasets of IDP conformations are obtained through all-atom MD simulations. Residues of IDP are encoded through a series of features, especially the frustration of residues is considered. GSALIDP can effectively predict interaction sites of IDP as well as the contact residue pairs between IDPs. Its performance in predicting dynamic IDP interaction is on par with or even better than the other models’ prediction of the conventional PPI. To the best of our knowledge, it is the first model to extend the PPI prediction to the IDP-involved interaction. Besides, the mechanism of these interactions and the relevance with residue frustration are discussed through the interpretability analysis of our model.

## Results

### Obtaining the conformation of IDPs from MD simulations

The N-terminal region of LAF-1, an intrinsically disordered R/G-rich domain (LAF-1 RGG domain), is considered in this work ^46,47^. Previous studies showed that this domain is important to the LLPS of LAF-1, which drives the assembly of the membraneless organelle called P granules ^46,48,49^. Besides, the isolated RGG domain can sufficiently engender LLPS *in vitro* ^24^. Thus, LAF-1 RGG domain is conceived as a representative IDP system ^15,24,46^.

In our previous work, all-atom MD simulations were conducted to characterize the interaction between two RGG domains ^15^. The trajectories are adopted for the current study. More specifically, 17 independent trajectories with a cumulative length of 18 μs are used. IDPs continuously undergo reversible interactions throughout the simulation. The durations of these interactions are highly varied. Apart from the transient interactions, there are stable interactions with sustained durations that dominate the IDP interaction ^15^. Accordingly, we collect a series of stable interaction events from the MD trajectories using the following threshold: two proteins keeping contact for more than 30 ns and the heavy atom contact number > 200. In total, 34 stable interaction events are obtained.

To consider the fluctuation of IDP’s flexible conformation, multiple IDP conformations prior to the interaction event are extracted. Specifically, 16 monomeric conformations of each IDP are obtained in 0.2 ns intervals between 0 to 3 ns prior to the contact. These conformations are used to calculate the features of the residues and construct the graphs that serve as the input of our model. The IDP complex at 3 ns after contact is used to label the interaction sites of each IDP as well as the contact residue pairs between two IDPs. The residues from two IDPs are considered to be in contact if the distance between any of their heavy atoms is less than 6 Å. The IDP complexes of the 34 interaction events are randomly divided into the training set and test set in the ratio of 6:1.

### Representation of IDP conformations

The conformation of IDP is represented as a graph G = (V, E), where V denotes the residues of IDP (nodes) and E denotes the contacts between residues (edges). This graph is encoded by a node feature matrix X and an adjacency matrix A: X is an n×d matrix that contains the feature values of the residues, where n is the number of residues and d is the number of features; A is an n×n matrix that encodes the contacts between residues, where the element A(i, j) equals 1 if there is a contact between residues i and j, and 0 otherwise.

For an IDP complex, 16 monomeric conformations of each IDP are considered (as described in the previous section). These conformations are represented as a sequence of graphs, {G_1_, G_2_, ⋯, G_16_}, where the G_1_, G_2_, …, G_16_ correspond to the conformation obtained at 3, 2.8, …, 0 ns prior to the contact, respectively. The n-th graph is denoted as G_n_ = (V_n_, E_n_), and its corresponding node feature matrix and adjacency matrix are denoted as X_n_ and A_n_. The prediction of interaction sites is based on the graph sequence of the IDP. And the prediction of the contact residue pairs is based on the graph sequences of both IDPs.

### Network architecture of GSALIDP

Multiple conformations of IDP (i.e. 16 conformations) are considered in our method to describe IDP’s flexibility. These conformations are represented by the sequence of graphs. In other words, IDP is characterized by a dynamic graph ^50–52^ instead of a single graph: the features of nodes (i.e., residues) and the connections between them (i.e., contacts) evolve over time. We develop the GraphSAGE-LSTM network that integrates the graph sample and aggregate (GraphSAGE) network ^53^ with the long short-term memory (LSTM) network ^54^ (see more discussions in Methods). LSTM serves as the main framework of the model, and GraphSAGE is embedded within LSTM to extract the structural information of IDPs. More specifically, GraphSAGE sequentially processes 16 IDP conformations and inputs the extracted structural information into the LSTM. LSTM further analyzes the hidden temporal information among these IDP conformations. This framework is similar to previous machine-learning methods of dynamic graph modeling ^50–52,55–57^, e.g. the integrated framework of graph convolutional network (GCN) and LSTM proposed by Seo et.al ^55^. The GraphSAGE-LSTM Network we develop can effectively extract the features of IDP’s conformational evolution.

The structure of our model is shown in Fig. 2. Standard LSTM comprises four core components: the input gate, forget gate, cell state, and output gate. In the GraphSAGE-LSTM network, the primary structure of LSTM is retained, but the computation of its four components is replaced by GraphSAGE. The GraphSAGE module generates node embeddings by sampling and aggregating features from a node’s neighborhood. Thus, the properties of the local environment of the node are effectively considered. And the architecture of LSTM can capture the long-term dependencies hidden in the sequence. The GraphSAGE-LSTM Network thus can effectively extract both the spatial and temporal information contained in the dynamic graph. It scans the entire graph sequence and outputs the hidden layer vector of the last graph. This vector is considered as the encoding of IDP residues and is further turned into the predicted results by a multi-layer perceptron (MLP). For the prediction of interaction sites, the MLP takes in the encoding of each residue and outputs the probability of its involvement in IDP interaction. For the prediction of contact residue pairs, the encodings of the residues of two IDPs are concatenated and the resulting vector is employed as the encoding of the residue pair between IDPs. The MLP then takes in this vector and outputs the contact probability of the residue pair.

**Fig. 1.**
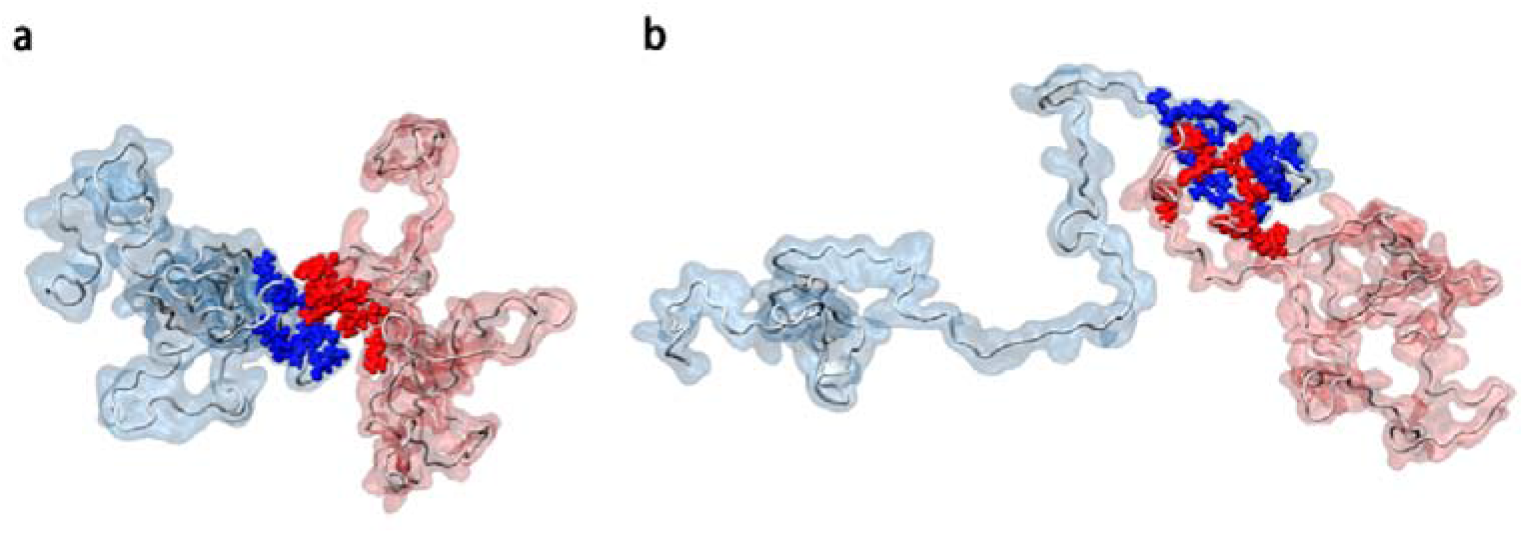
Representative complex structures of LAF-1 RGG domains obtained from simulations. In red and blue are represented two proteins and in spheres are represented the residues involved in IDP interaction.

**Fig. 2.**
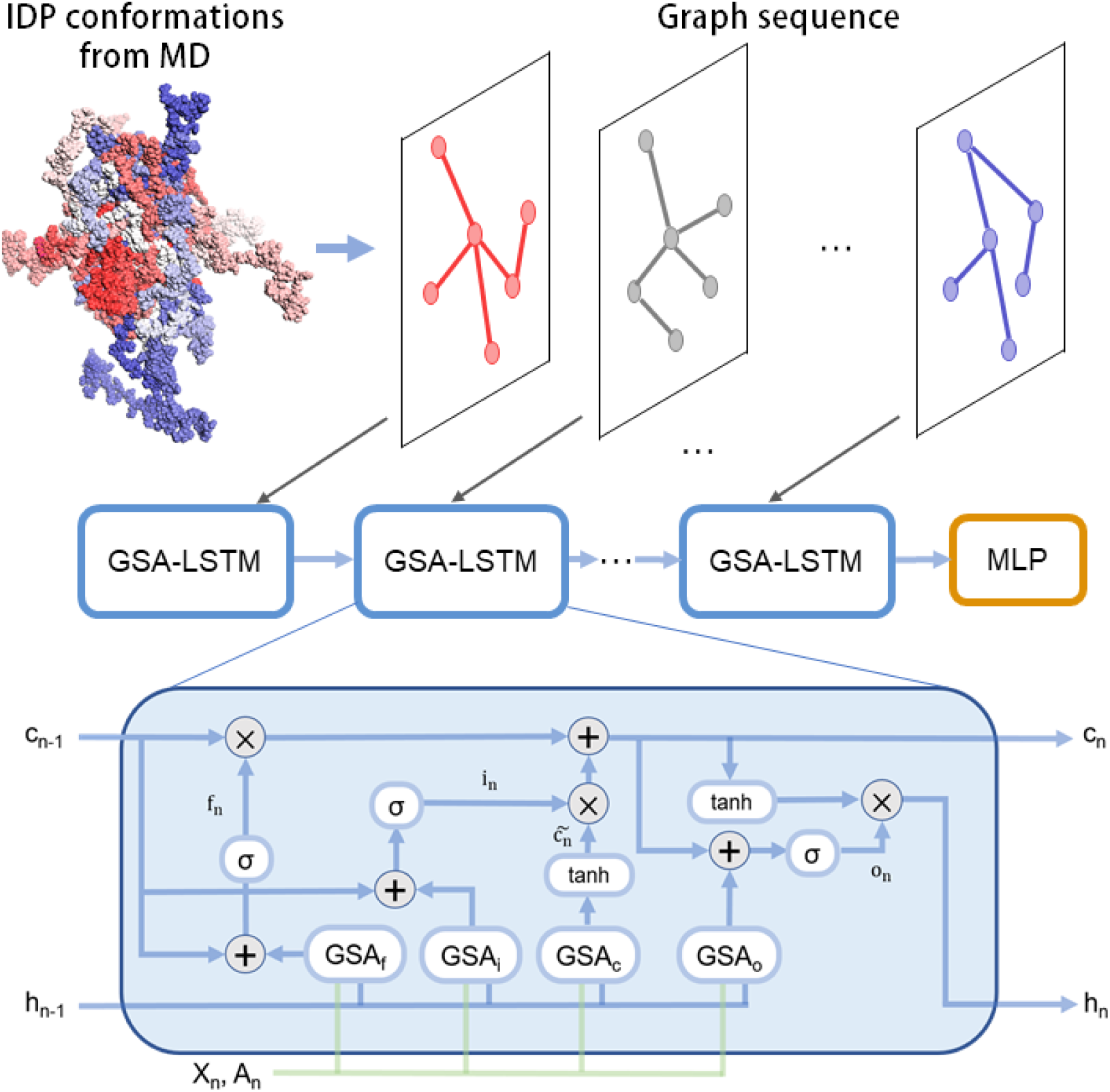
The overall framework of GSALIDP. The input of GSALIDP is a sequence of graphs {G_1_, …, G_N_} where each graph represents an IDP conformation and is encoded by the node feature matrix X_n_ and the residue contact matrix A_n_. The GraphSAGE-LSTM Network (GSA-LSTM) scans the entire graph sequence and outputs the hidden layer vector of the last graph. A multi-layer perceptron (MLP) then converts this vector into the predicted probabilities. Here, h_n_ and c_n_ respectively represent the hidden layer vector and cell state. GSA_f_, GSA_i_, GSA_c_, and GSA_o_ represent the GraphSAGE-embedded gates of the LSTM framework.

### Performance of GSALIDP in predicting the interaction sites of IDP

The receiver operating characteristic (ROC) curve shows the pairs of true positive rate and false positive rate calculated at all possible thresholds. The ROC curve for the model with a random performance is a straight diagonal line from (0, 0) to (1, 1) and the corresponding area under the curve (AUROC) is 0.5 ^58^. It serves as the baseline of the ROC curve. The ROC curve of GSALIDP’s prediction on the interaction sites is represented by the yellow curve in Fig. 3. This curve exceeds the baseline and the corresponding AUROC is up to 0.81. Thus, our model can effectively predict the interaction site between IDPs. And the performance is even better than the mainstream PPI prediction methods of structural proteins ^32–40^, whose AUROCs are in the range of 0.58 to 0.79 ^40^.

**Fig. 3.**
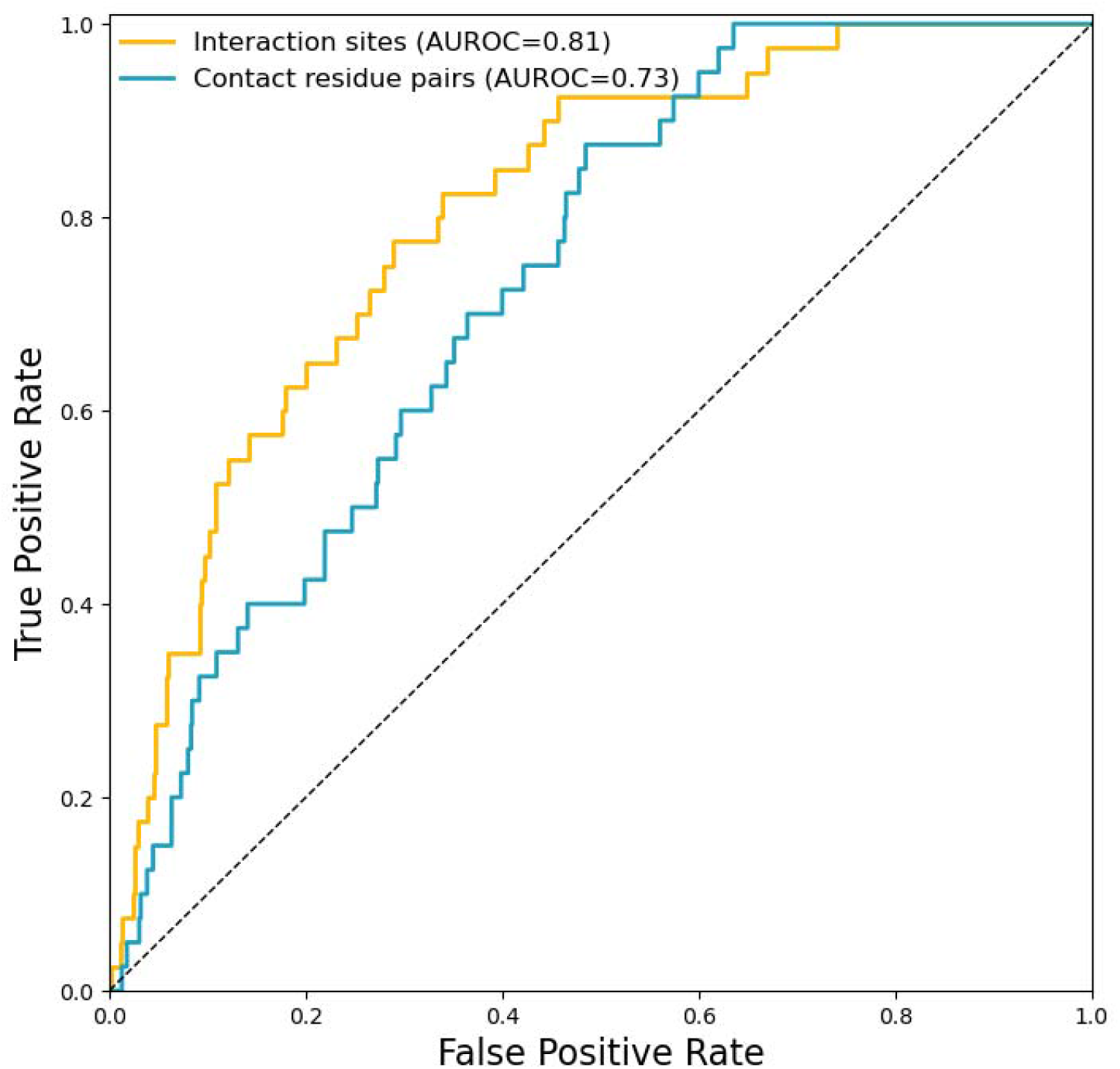
ROC Curves of GSALIDP. The curves for interaction sites, contact residue pairs, as well as random predictions are shown by the yellow solid line, blue solid line, and black dotted line, respectively.

In addition to AUROC, we also consider the area under the precision-recall curve (AUPRC), F1-score (F1), and Matthews correlation coefficient (MCC). These metrics can also effectively measure the performance of PPI prediction models^32–34,40^. Other metrics including accuracy (ACC), precision, and recall are also used. For all metrics we considered (Table 1), our model achieves comparable performance with the conventional PPI prediction models of structural protein ^32–40^. Our model outperforms some mainstream models of structural protein with respect to metrics like AUROC, MCC, etc. It should be noted that the available data for IDP interactions is highly scarce as compared to the structural protein counterpart. Moreover, the dynamic nature of IDP interactions provides additional challenges to the prediction. These results further demonstrate GSALIDP’s ability to distinguish between the interaction and non-interaction sites of IDPs.

**Table 1.** Performance of interaction site prediction.

| Method | AUROC | AUPRC | MCC | F1 | Precision | Recall | ACC |
| --- | --- | --- | --- | --- | --- | --- | --- |
| Conventional PPI models | 0.58 ~ 0.79 | 0.19 ~ 0.44 | 0.08 ~ 0.34 | 0.28 ~ 0.46 | 0.19 ~ 0.37 | 0.41 ~ 0.59 | 0.57 ~ 0.78 |
| GSALIDP | 0.81 | 0.16 | 0.26 | 0.28 | 0.19 | 0.53 | 0.87 |

*Note:* Performance of the prediction models of structural proteins are obtained from Yuan et al <sup>40</sup>.

### Performance of GSALIDP in predicting the contact residue pairs between IDPs

It should be noted that the stability of IDP interaction is attributed to the formation of contact residue pairs^15^. Under such consideration, the contact residue pairs between IDPs are also predicted. The ROC curve of GSALIDP on predicting the contact residue pairs is represented in Fig.3. The AUROC is up to 0.73. Such performance is also comparable with previous models for predicting the contact residue pairs between structural proteins^41–44^, whose AUROCs are in the range of 0.69 to 0.91 ^43,44^.

### Interpretations of our model

The impressive performance of GSALIDP indicates it can effectively predict the behavior of IDP interaction. Previous studies highlighted the importance of interpretability analysis of machine-learning models, which can facilitate obtaining new insights and extracting the knowledge that was originally hidden in the data ^59–61^. Thus, we discuss the possible mechanism underlying IDP interaction by evaluating the relative importance of the utilized features.

The residues of IDP are characterized by a set of features (see Methods). Specifically, both the conventional geometric/physicochemical-based features and the statistics-based features related to the frustrations of residue interactions are considered. To demonstrate the relative importance of these features, a strategy for interpreting machine-learning models called sensitivity analysis is conducted ^61–65^.

Sensitivity analysis involves excluding one feature from the feature set and measuring the impact on the model’s performance. Supplementary Table 2 summarizes the sensitivity analysis of all features used in our model. Among these features, the removal of the relative highly frustrated density (RHFD) or the interior contact area (ICA) has the most significant effect. The performance of our model after the exclusion of RHFD and ICA as well as only considering the RHFD and ICA is compared (Table 2). The exclusion of either RHFD or ICA results in considerable declines in the model’s AUROC, AUPRC, F1, and MCC. Moreover, if the feature set comprises only RHFD and ICA, the model can still realize an acceptable predictive performance. Taken together, RHFD and ICA are important for predicting IDP interaction. Additional discussions with regard to these two features are then conducted as follows.

**Table 2.** Performance of interaction site prediction using different feature sets.

| Features | AUROC | AUPRC | F1 | MCC | Precision | Recall | ACC |
| --- | --- | --- | --- | --- | --- | --- | --- |
| All features | 0.81 | 0.16 | 0.26 | 0.28 | 0.19 | 0.53 | 0.87 |
| Exclude RHFD | 0.66 | 0.09 | 0.18 | 0.14 | 0.12 | 0.38 | 0.84 |
| Exclude ICA | 0.69 | 0.11 | 0.17 | 0.17 | 0.10 | 0.70 | 0.68 |
| Using only ICA and RHFD | 0.68 | 0.10 | 0.14 | 0.13 | 0.08 | 0.73 | 0.59 |

The relative highly frustrated density (RHFD) of a residue is defined as the ratio between the number of highly frustrated interactions and the number of all interactions that the residue forms ^44^. Residues with higher RHFDs tend to encounter more severe local energy conflicts. Residues are categorized into groups according to their RHFD values, and the percentage of interaction sites in each group is calculated (Fig. 4a). It should be noted that the overall percentage of residues with high RHFD (i.e. between 0.6 and 1) is very low (Supplementary Fig. 2), these residues are combined into one group. With the increase of the RHFD, the percentage of interaction sites also increases, suggesting a clear correlation between the residue’s RHFD and its tendency to participate in PPI. Such correlation with PPI involvement may be attributed to the fact that local energy conflicts within IDP monomer can be reduced by the interactions with other IDP. As revealed in a recent study, frustration in the complex is lower than in the free-state monomer ^66^. The RHFD of residues in monomeric IDP is then compared with the cases of IDP complex. The average RHFD of the interaction sites in the monomeric IDP is 0.131. Interestingly, the average RHFD of interaction sites within the IDP complex significantly decreases to 0.076. Such observation corroborates our conjecture and illustrates the importance of the energy conflicts within IDP.

**Fig. 4.**
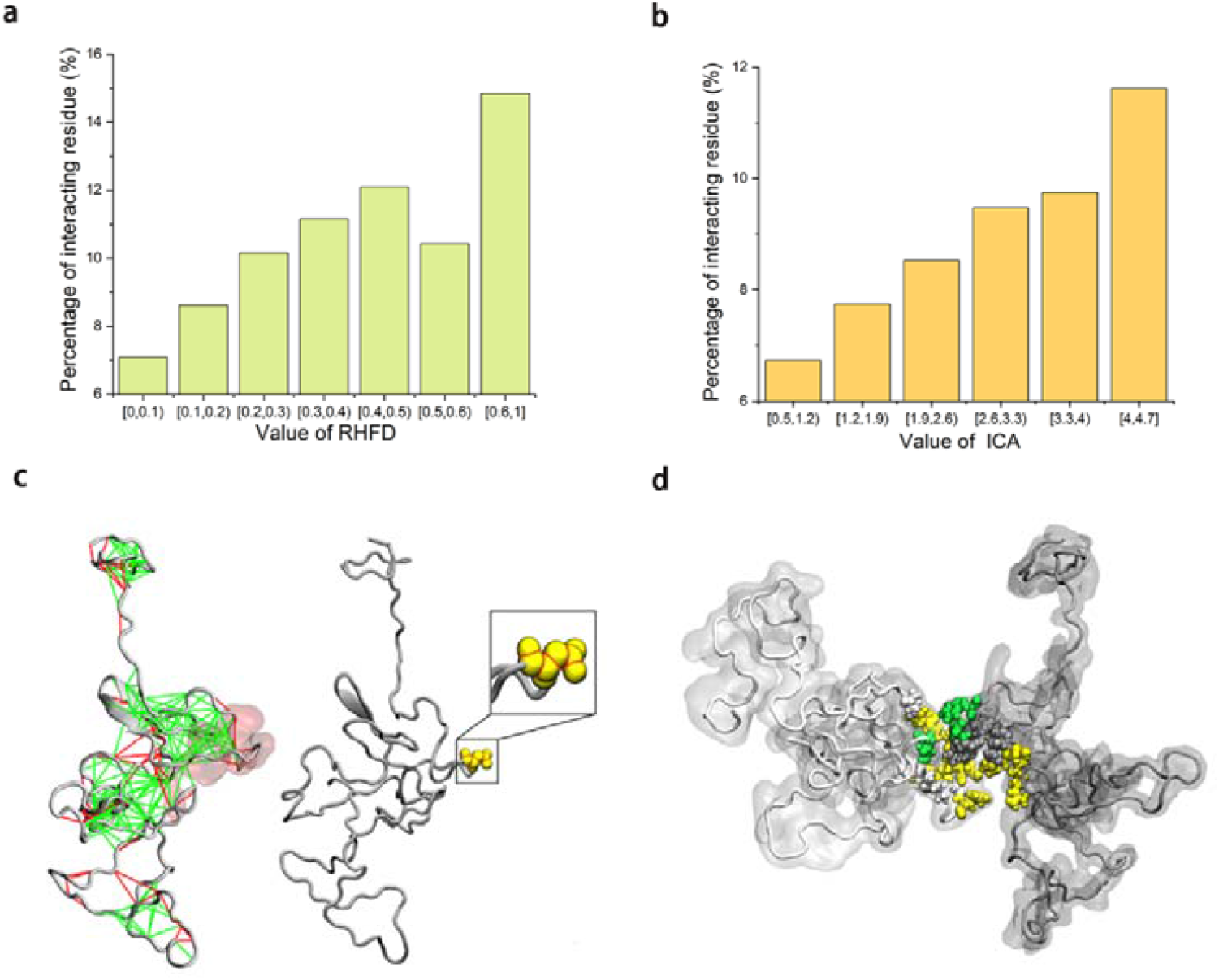
Analysis of ICA and RHFD. (a-b) Percentage of the interaction sites in the residue groups with different RHFD or ICA. (c) Left, frustratogram of the representative IDP conformation highlighted the highly (red) and minimally (green) frustrated interactions, and PPI residues (red surface). Right, diagram of ICA of a residue. The red line represents the outline of the ICA. (d) Rrepresentative conformation of IDP complex with highlighted interaction sites. The residues with high RHFD or high ICA are shown in green and yellow, respectively.

The interior contact area (ICA) is defined as the sum of the contact areas of all atoms within the residue and is calculated using the Qcontacts package ^67^. We group the residues according to their ICA values and calculate the percentage of interaction sites in each group. As the ICA increases, the proportion of interaction sites also increases considerably (Fig. 4b). Residues with higher ICA values have more contracted side chains and their backbones are more exposed. Such conformations should favor the backbones to participate in the IDP interaction. Thus, the observed correlation may be attributed to the importance of backbone-involved interactions. According to our analysis of all complexes in the dataset, there are 62 % interaction residues whose backbone moieties participate in the interaction with the residues of the other IDP.

As demonstrated by these results, the extent of frustration and the backbone exposure of the residues are directly correlated to their tendencies to participating IDP interaction. Accordingly, for a given IDP conformation there are certain regions that have apparent propensities to mediate stable interaction with other IDP, i.e. the regions enriched of the residues with these features. Fig. 4d and Supplementary Fig. 3 illustrate representative conformations of IDP complex with highlighted interaction sites. The residues with high RHFD or high ICA (top one third of residues) are shown in green and yellow, respectively. There is an apparent enrichment of the residues with high RHFD or high ICA. We further analyze the complexes throughout the dataset, and there are more than 61 % of interaction sites with high RHFD or high ICA. The enrichment of the residues with these features in the binding interface also demonstrates the importance of RHFD and ICA.

## Discussion

In this work, we present GSALIDP, a novel GraphSAGE-LSTM Network for predicting the behavior of IDP-involved interactions. This framework models multiple conformations of IDP as a dynamic graph to describe the considerable fluctuation of IDP’s flexible conformation. Its performance in predicting the dynamic IDP interaction is on par with or even better than the other models’ prediction of conventional PPIs. Besides, the interpretability analysis of GSALIDP identifies two important features for the prediction of IDP interaction, i.e. RHFD and ICA, which reveal the importance of the energy conflicts and backbone moiety. As shown by our work, the dynamic nature of IDP interaction can be effectively depicted by the dynamic graph neural network. This model can reliably predict the interaction between IDPs. This framework can be exploited for the studies of other IDP-involved interactions.

## Materials and Methods

### Features of the residue

The residues of IDP are characterized by a series of features. Here, statistics-based features, geometric-based features, and physicochemical-based features are all considered.

#### Statistics-based features

As demonstrated in our previous work, statistics-based indexes related to the frustrations of residue interaction are effective features for PPI prediction ^44^. The frustration index at contact level is calculated for each residue-residue contact present in a protein structure, which is a statistical measure of how energetically favorable a native contact is with respect to a set of all possible contacts at that location. A recent study showed that local frustration is an evolutionary constraint that is related to protein stability and function ^68^. As indicated in previous studies, residues involved in PPIs are enriched with highly frustrated contacts ^69–72^, suggesting the frustration index may be a suitable feature for PPI prediction. Thus, the features based on the frustration index, i.e. the highly frustrated densities (HFD) and the relative highly frustrated density (RHFD), are applied to encode the residues.

The HFD and the RHFD are calculated using the Frustratometer tool ^73,74^. This algorithm quantifies the frustration in the protein structure obtained by our simulation. The frustration index (FI) can be calculated in two modes at the contact level, i.e. mutational and configurational FI. In this work, the configurational FI is considered. It is calculated based on the decoys that change both the identity of the amino acids in contact and the contact distance accordingly. Thus, configuration FI can effectively characterize the frustrations of residue contact within the flexible structure. The FI value is calculated as a z-score of the distribution of the difference between the native energy of the protein and the energy of the decoys. The FI can be classified according to its value into minimally frustrated (FI > 0.78), neutral (−1 < FI < 0.78), and highly frustrated (FI < −1). After classifying the contacts, the HFD and the RHFD of the residues can be obtained. The residue’s HFD is the number of involved highly frustrated interactions, while the residue’s RHFD is the HFD divided by the number of all the involved interactions.

#### Geometric-based and physicochemical-based features

The commonly used geometric-based features like solvent accessible surface area (SASA) and relative solvent accessible surface area (RASA) as well as the lately developed features including interior contact area (ICA), exterior contact area (ECA), and exterior void area (EVA) are used to describe residues ^75–78^. In addition, physicochemical-based features are also applied, including the hydropathy index (HI, two versions) and pK_α_ (two versions) ^79,80^. The calculation methods of these features are described in supplementary materials.

### The GraphSAGE-LSTM Network

For each interaction event, multiple conformations of IDP are considered to describe the fluctuation of IDP’s flexible conformation. These conformations are considered as a dynamic graph, where the features of the nodes (i.e., residues) and the connections between them (i.e., contacts) evolve dynamically over time. Similar dynamic graphs have been used to model a variety of time-evolving networks, such as social networks, wireless networks, and others ^51,52,55^. A series of machine-learning methods have been developed for embedding these dynamic graphs^50–52,55–57^. Especially, the framework that combines graph neural network (GNN) and recurrent neural network (RNN) is leveraged to capture both the spatial and temporal features of the dynamic graph _50,55,57_.

Inspired by these studies, we develop the GraphSAGE-LSTM network. The main framework of the LSTM network is retained, but the computations of its input gate, forget gate, cell state, and output gate are replaced by GraphSAGE networks. The GraphSAGE-LSTM network sequentially processes the IDP conformations in the graph sequence {G_1_, …, G_N_} and outputs the hidden state h_N_ of the last graph, which serves as the embedding of the residues. For clarity, the processing of the n-th graph (where n=1,…, N) is described as follows:

First, the input gate i_n_, forget gate f_n_, and output gate o_n_ are determined using:

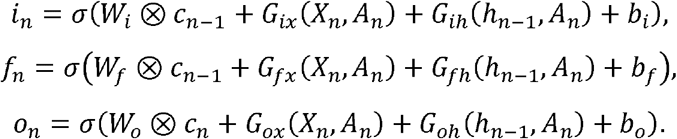

c_n-1_ and h_n-1_ denote the cell state and hidden state of the preceding graph G_n-1_, respectively. For the initial conformation (n=1), c_0_ and h_0_ are set as zero tensors. X_n_ represents the tensor of residue features and A_n_ is the adjacency matrix of the n-th graph. *G* represents the GraphSAGE network^53^. W_i,f,o_, b_i,f,o_ denote the trainable parameters of the LSTM network.

After the calculations of the i_n_, f_n_, and o_n_, the cell state c_n_ and hidden state h_n_ of the n-th graph are calculated using:

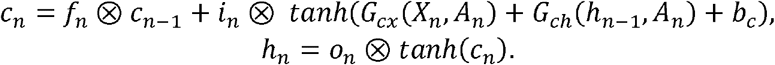

The c_n_ and h_n_ serve as the output of the n-th graph, and will be fed into the model for processing the subsequent graph. After scanning the entire sequence, the model outputs the hidden state h_N_ of the last graph. This h_N_ serves as the embedding of the residues. The dimension of h_N_ is n_r_×d_h_, where n_r_ and d_h_ represent the number of residues and the hidden layer width, respectively.

A multilayer perceptron (MLP) consisting of four linear layers followed by a softmax layer is then utilized to output the prediction results. For the interaction site prediction, the MLP receives the embedding of the residues (h_N_). The first three linear layers successively halve the dimension of the hidden layer width d_h_, and the final linear layer reduces the dimension to 2 and outputs a tensor with a dimension of n_r_×2. The softmax layer then normalizes this tensor into a probability distribution over the two predicted classes (non-interacting and interacting). For the contact residue pair prediction, the MLP receives the embedding of the residue pairs between IDPs, which is obtained by concatenating the embeddings of the residues of IDPs. The linear layers successively reduce the dimension of the hidden layer width and output a tensor with a dimension of n_r_^2^×1. The softmax layer then converts this tensor into the predicted contact probabilities of the residue pairs.

### Evaluation metrics

The model’s performance is evaluated by a series of metrics. The principle metric we considered is the area under the receiver operating characteristic curve (AUROC). AUROC is the area under the ROC curve that shows pairs of true positive rate and false positive rate values calculated at all possible thresholds ^58^. AUROC reveals the overall performance of a classifier: 0.5 is for random classifiers and 1.0 is for perfect classifiers. Thus, a model with an AUROC higher than 0.5 is considered to make effective predictions^58^. Similar to previous PPI prediction works of structural proteins, other metrics are also considered. Specifically, the area under the precision-recall curve (AUPRC), F1-score (F1), and Matthews correlation coefficient (MCC) are employed. These metrics are capable of effectively measuring the performance of PPI prediction models ^32–34,40^. In addition, other metrics including accuracy (ACC), precision, and recall are also employed. The AUROC and AUPRC are independent of the thresholds. The other metrics are calculated using a threshold to convert predicted probabilities to binary predictions. And the threshold is determined by maximizing the F1-score of the model. The formulas for calculating these metrics are provided in the supplementary materials.

### Implementation details

For the interaction site prediction, we perform the leave-one-out cross validation on the training set. Each complex takes turns to evaluate the model trained on the rest of the complexes in the training set. The resulting performances are then averaged as the overall validation performance, which is adopted to choose the optimal hyperparameters and training epoch. The chosen hyperparameters are as follows: hidden size = 128, learning rate = 1e-3, dropout rate = 0.3, weight decay of MLP = 1e-2, and weight decay of the GraphSAGE-LSTM network = 5e-4. The model is then retrained on the entire training set with the optimal hyperparameters and training epoch and is evaluated on the independent test set. For the contact residue pair prediction, we perform 10-fold cross validation. The training set is randomly split into 10 folds. Each time, a model is trained on nine folds and evaluated on the remaining one fold. This process is repeated ten times and the performances are averaged to obtain the overall validation performance. And the chosen hyperparameters are as follows: hidden size = 128, learning rate = 1e-3, dropout rate = 0.3, weight decay of MLP = 1e-2, and weight decay of the GraphSAGE-LSTM network = 5e-4. The model is then retrained on the entire training set and evaluated on the test set.

We implemented the proposed model with Pytorch 1.12.1 ^81^, PyTorch Geometric 2.3.0 ^82^, and PyTorch Geometric Temporal 0.54.0 ^83^. And the Cross-Entropy loss with weights and Adam optimizer are employed for optimization ^84^.

## Acknowledgments

We thank Prof. Rui Shi and Prof. Qianyuan Tang for their comments on the manuscript.

## Author Contributions

J-Y. L. designed and guided the study. Y-C. Z. prepared the dataset, designed the model architecture, and wrote the first draft. Q-X. L. designed the model architecture and implemented the method. All authors discussed the results and contributed to the writing of the manuscript.

## Data and code availability

The IDP conformations and the processed feature values are available on GitHub (https://github.com/arnoldland/GSALIDP). The code of the proposed model is available on GitHub (https://github.com/arnoldland/GSALIDP).

