## Supplementary Information for "Predicting the dynamic interaction of intrinsically disordered proteins" (from another version of this preprint: version 1, 2023-12-04)

### Supplementary Methods

**Calculation of evaluation metrics.** A series of evaluation metrics are used to evaluate our model's predictive performance. The accuracy (ACC), precision, recall, F1-score (F1), and Matthews correlation coefficient (MCC) are calculated using:

$$\mathbf{ACC} = \frac{\boldsymbol{TP+TN}}{\boldsymbol{TP+TN+FP+FN}}$$

$$\mathbf{Precision} = \frac{\boldsymbol{TP}}{\boldsymbol{TP+FP}}$$

$$\mathbf{Recall} = \frac{\boldsymbol{TP}}{\boldsymbol{TP+FN}}$$

$$\mathbf{F1} = \mathbf{2} \times \frac{\boldsymbol{Precision} \times \mathbf{Recall}}{\boldsymbol{Precision} + \mathbf{Recall}}$$

$$\mathbf{MCC} = \frac{\boldsymbol{TP} \times \mathbf{TN} - \mathbf{FN} \times \mathbf{FP}}{\sqrt{(\boldsymbol{TP+FP}) \times (\mathbf{TP+FN}) \times (\mathbf{TN+FP}) \times (\mathbf{TN+FN})}}$$

where true positives (TP) and true negatives (TN) denote the number of correctly identified interacting and non-interacting sites, and false positives (FP) and false negatives (FN) denote the number of incorrectly identified interacting and non-interacting sites, respectively. These metrics are calculated using a threshold to convert predicted interacting probabilities to binary predictions. The threshold is determined by maximizing the F1-score.

AUROC and AUPRC are independent of thresholds. AUPRC is the area under the precision-recall curve which shows pairs of precision and recall values. AUROC is the area under the ROC curve that shows pairs of true positive rate (TPR) and false positive rate (FPR) values, where the TPR and FPR are calculated using:

$$\mathbf{TPR} = \frac{\boldsymbol{TP}}{\boldsymbol{TP+FN}}$$

$$\mathbf{FPR} = \frac{\boldsymbol{FP}}{\boldsymbol{TN+FP}}$$

**Calculation of geometric-based and physicochemical-based features.** The commonly used geometric-based features like solvent accessible surface area (SASA) and relative solvent accessible surface area (RASA) as well as the lately developed features including interior contact area (ICA), exterior contact area (ECA), and exterior void area (EVA) are used to describe residues[1–4]. SASA is calculated using the GROMACS package[5]. The calculation algorithm is the rolling ball algorithm, and the radius of the probe is 0.14 nm. RASA is calculated by the following formula:

$$\mathbf{RASA} = \frac{\boldsymbol{SASA}_{\boldsymbol{in\ chain}}}{\boldsymbol{SASA}_{\boldsymbol{unbound}}}$$

where $SASA_{in\ chain}$ refers to the SASA of the residue in the protein monomer, and $SASA_{unbound}$ residue refers to the SASA of the isolated residue.

ICA and ECA are calculated using the Qcontacts package with default parameters [6]. EVA is calculated using:

$$\mathbf{EVA} = \mathbf{SASA}_{\mathbf{unbound}} - \mathbf{SASA}_{\mathbf{in\ chain}} - \mathbf{ECA}$$

Physicochemical-based features including two versions of hydropath index (H1 and H2) and two versions of pKa (pKa1 and pKa2) are also considered. pKa1 and pKa2 are calculated using the PROPKA3 package[7]. And H1 and H2 are obtained from references [8,9].

### Supplementary Tables

**Supplementary Table 1 |** AUROC of the models with different feature sets.

| Excluded features | SASA | RASA | ICA | ECA | EVA | HFD |
|---|---|---|---|---|---|---|
| AUROC | 0.82 | 0.75 | **0.69** | 0.78 | 0.71 | 0.76 |
| Excluded features | RHFD | HI_1 | HI_2 | pKα_1 | pKα_2 | |
| AUROC | **0.66** | 0.73 | 0.78 | 0.75 | 0.79 | |

### Supplementary Figures

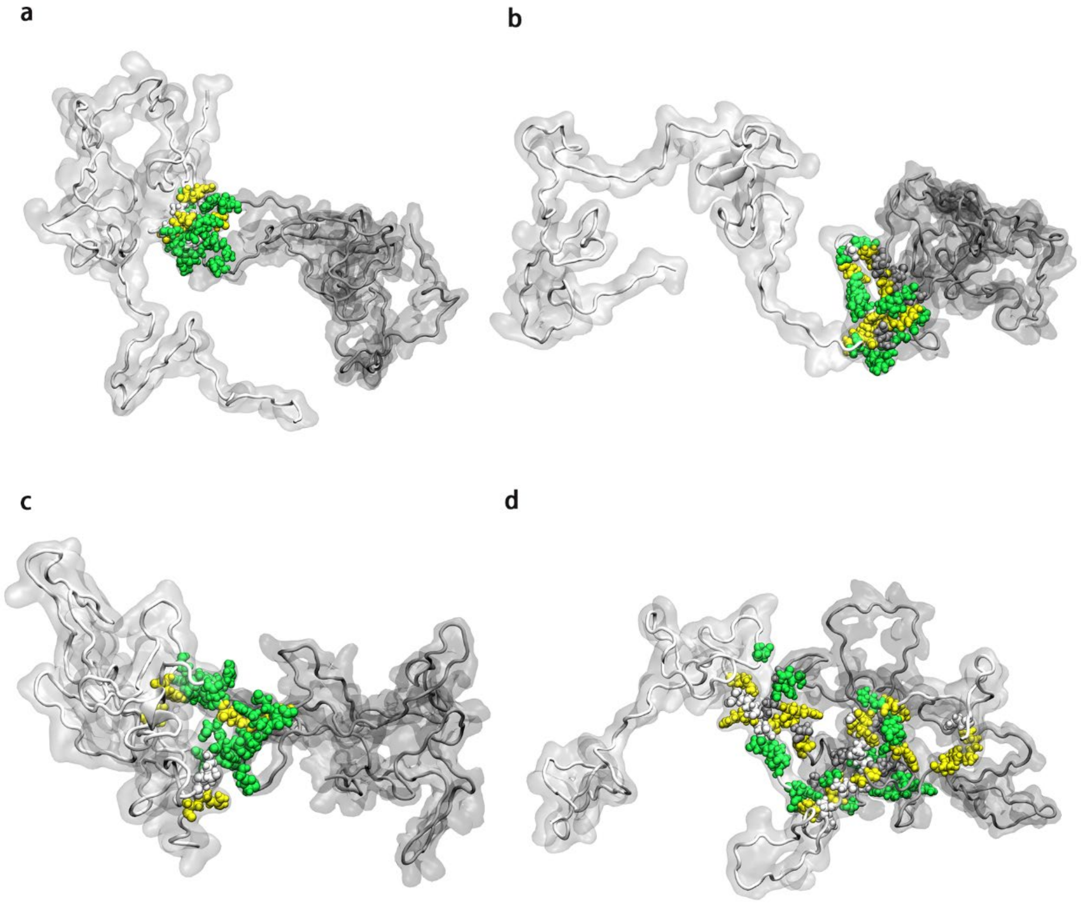


**Supplementary Figure 1 |** Rrepresentative conformations of IDP complex with highlighted interaction sites. The residues with high RHFD or high ICA are shown in green and yellow, respectively.

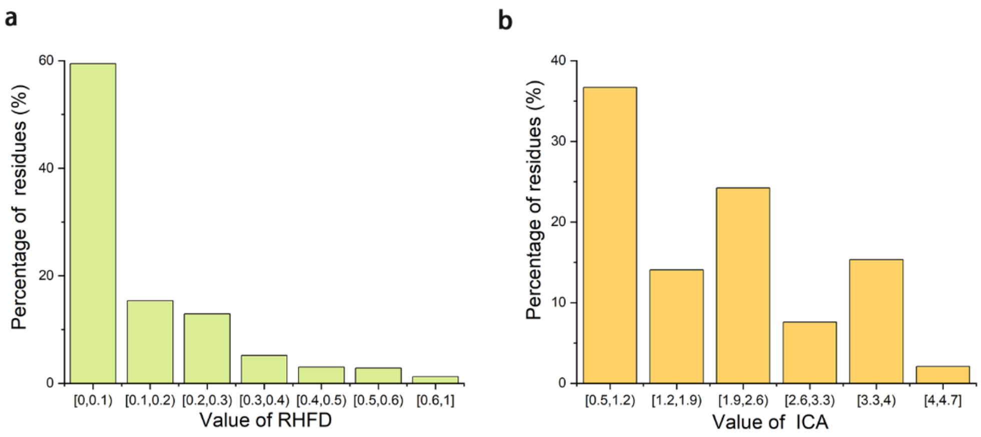


**Supplementary Figure 2 | a-b,** Overall percentage of the residues in the groups with different RHFD or ICA.

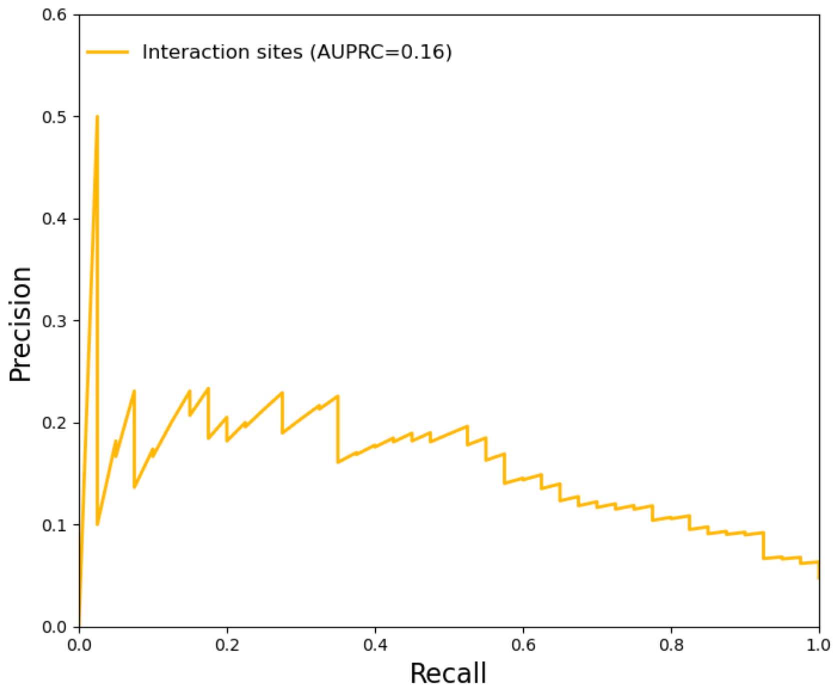


**Supplementary Figure 3 |** Precison-recall curve (PRC) of GSALIDP on test set.
